# The B-box protein BBX13/COL15 suppresses photoperiodic flowering by attenuating the action of CONSTANS in Arabidopsis

**DOI:** 10.1101/2024.04.19.590210

**Authors:** Puthan Valappil Rahul, Premachandran Yadukrishnan, Anagha Sasidharan, Sourav Datta

## Abstract

The optimal timing of transition from vegetative to floral reproductive phase is critical for plant productivity and agricultural yields. Light plays a decisive role in regulating this transition. The B-box (BBX) family of transcription factors regulates several light-mediated developmental processes in plants, including flowering. Here, we identify a previously uncharacterized group II member of the BBX family, BBX13/COL15, as a negative regulator of flowering under long-day conditions. *BBX13* is primarily expressed in the leaf vasculature, buds, and flowers. Its spatial expression pattern is similar to the major flowering time regulators *CO* and *FT. bbx13* mutants flower early, while *BBX13*-overexpressors exhibit delayed flowering under long days. Genetic analyses showed that *BBX13* acts upstream to *CO* and *FT* in the flowering pathway and negatively regulates their expression. We found that BBX13 physically interacts with CO and inhibits the CO-mediated transcriptional activation of *FT*. In addition, BBX13 directly binds to the *CORE2* motif on the *FT* promoter, a site where CO also binds. Furthermore, our ChIP data indicates that BBX13 reduces the in vivo binding of CO on the *FT* promoter. Through luciferase assay, we found that BBX13 inhibits the CO-mediated transcriptional activation of *FT.* All these data together suggest that BBX13/COL15 represses flowering in Arabidopsis by attenuating CO activity.

**Summary statement:** BBX13, a previously uncharacterized protein in the B-Box family of transcription factors, negatively regulates flowering time in *Arabidopsis thaliana*. BBX13 physically and genetically interacts with CO and inhibits the CO-mediated transcriptional activation of *FT*.

## Introduction

Plants must integrate many endogenous and exogenous signals to mediate their growth and development. Light is one of the most important exogenous signals that plants perceive. Plants possess several proteins that mediate light-signaling pathways and photo-responses (Srikanth and Schmid, 2011). In plants, flowering is one of the most important developmental switches that determines its reproductive success. Flowering is controlled by different signaling pathways that converge to create a robust seasonal response (Andrés and Coupland, 2012). The major flowering regulatory pathways are photoperiodic, thermo-sensory, autonomous, vernalization, gibberellic acid-mediated, and age-related (Gursky et al., 2018; Takagi et al., 2023). Light is an essential environmental factor that regulate flowering time through its duration (photoperiod), quality and intensity. Different plants are classified as long-day, short-day, or day-neutral, based on the differences in the length of uninterrupted dark periods they require to initiate flowering. *Arabidopsis thaliana* is a facultative long-day plant that flower early in the inductive photoperiods (16h light/ 8h dark) (Luo et al., 2021).

The B-box (BBX) family of proteins, which are zinc finger transcription factors, is known to play versatile roles in light-regulated development (Song et al., 2020; Yadav et al., 2020a; Cao et al., 2023). B-box family members contain one or more B-box domains comprised of highly conserved repetitive sequences of cysteine and histidine, with specialized tertiary structures that are stabilized by binding to Zn ions (Klug and Schwabe, 1995). Sometimes, they also feature a CCT (CONSTANS, CO-like, and TOC1) domain towards their C-terminal region. They regulate various light-regulated developmental pathways including seed germination, photomorphogenesis, cotyledon opening and greening, anthocyanin accumulation, shade avoidance responses, circadian clock, and flowering (Khanna et al., 2009; Gangappa and Botto, 2014; Vaishak et al., 2019; Song et al., 2020; Yadav et al., 2020a; Job and Datta, 2021; Ravindran et al., 2021).

BBX1/CONSTANS (CO), the first member of the BBX family identified in Arabidopsis, belongs to structural group I and contains two B-box motifs and one CCT domain. Light and the circadian clock coordinate CO/BBX1 activity to promote flowering under long-day photoperiods (LDs - 16h light/ 8h dark) in Arabidopsis (Putterill et al., 1995; Valverde et al., 2004). Under LDs, CO activates the transcription of flowering inducers such as *FLOWERING LOCUS T (FT)* and *SUPPRESSOR OF OVEREXPRESSION OF CONSTANS1 (SOC1)* (Samach et al., 2000; Hayama et al., 2003; Abe et al., 2005; Wigge et al., 2005; Yoo et al., 2005; Searle et al., 2006). The circadian clock mediates the transcriptional regulation of CO, while the turnover of CO protein is regulated by E3 ubiquitin ligases like CONSTITUTIVELY PHOTOMORPHOGENIC 1 (COP1) and HIGH EXPRESSION OF OSMOTICALLY RESPONSIVE GENES 1 (HOS1) (Jang et al., 2008; Lazaro et al., 2012). COP1 dimerization occurs during darkness and dawn, degrading the CO protein by its E3-ubiquitin ligase activity. FKF1 inhibits this dimerization during the evenings of LDs, which in turn induces the protein levels of CO (Lee et al., 2017). CO binds to the CO-Responsive Elements-*CORE1* and *CORE2* motifs of the *FT* promoter and positively regulates the transcription of *FT* (Tiwari et al., 2010). FT protein translocates from the leaves to the shoot apex. It forms a complex with the basic leucine zipper (bZIP) transcription factor FLOWERING LOCUS D (FD) to activate the expression of the floral meristem identity genes such as *APETALA1* (*AP1*), *LEAFY* (*LFY*), and *FRUITFULL* (*FUL*) (Abe et al., 2005; Wigge et al., 2005; Jung et al., 2016). While CO acts as a primary determinant of the transition to the reproductive phase, several proteins act on the CO-FT module to fine-tune the flowering time (Srikanth and Schmid, 2011). For example, other members of the BBX family positively and negatively regulate flowering time through multiple mechanisms such as the direct regulation of *FT* transcription, transcriptional regulation of *CO*, and modulation of the activity of CO via protein-protein interaction (Cheng and Wang, 2005; Datta et al., 2006; Hassidim et al., 2009; Li et al., 2014; Wang et al., 2014; Graeff et al., 2016; Tripathi et al., 2016; Ordoñez-Herrera et al., 2018; Steinbach, 2019; Liu et al., 2020; Wang et al., 2021; Xu et al., 2022). In addition, multiple subunits of NUCLEAR FACTOR-Y proteins (NF-Ys) interact with CO and mediate the binding of CO on the *FT* promoter through CCAAT motifs to regulate flowering time (Wenkel et al., 2006; Zhao et al., 2017).

Although extensive studies in the last two decades have led to the characterization of most of the proteins in the thirty-two-membered BBX family in Arabidopsis, the physiological role of a rare few remains to be understood. In this study, we explored the role of a previously uncharacterized BBX family transcription factor, BBX13, previously known as COL15 (CONSTANS-LIKE15), and showed that BBX13/COL15 negatively regulates flowering time in Arabidopsis by inhibiting CO-mediated transcriptional activation of *FT*.

## Materials and Methods

### Plant materials and growth conditions

*Arabidopsis thaliana* ecotype Col-0 (Columbia-0) was used as the wild type for all experiments in this study. The *bbx13-1* (SALK_074832) *and bbx13-2* (SALK_106851) mutants were obtained from ABRC, and the homozygous lines were identified by PCR-based genotyping. Sequences of all the primers used in the study are listed in the **Table S1**. *co-SAIL* and *ft-10* were previously described (Yoo et al., 2005; Laubinger et al., 2006). *35S:CO-YFP* was generously provided by Dr. Utpal Nath (Kumar et al., 2018). Seeds were surface-sterilized with 4% bleach for 3 min and washed with water five times before being sown on plates with Murashige and Skoog medium supplemented with sucrose and 0.8% agar. After stratification for 3 days in the dark at 4°C, the plates were transferred to Percival growth chambers with a fluence of ∼100 μmol m^−2^ s^−1^ and an ambient temperature of 22°C under the specified light conditions. Long-day conditions (LDs) were maintained at 16h light/8h dark while the short-day conditions (SDs) were at 8h light/16h dark. The *co bbx13-1*and *ft-10 bbx13-1* double mutants were generated by crossing *bbx13-1* with the respective single mutants.

### Generation of transgenic lines

The *BBX13* full-length coding sequence was amplified from a Col-0 cDNA library using Gateway-compatible primers (**Table S1**) and was cloned into the entry vector *pDONR207* using the BP reaction. The entry vector was used to perform an LR reaction with a modified *pCAMBIA1300* to generate the *35S:BBX13* construct. After sequencing, the plasmid was transformed into Agrobacterium strain GV3101, and cultures of the confirmed colonies were used for the floral dipping method of transformation in Col-0 (Clough and Bent, 1998; Zhang et al., 2006). Transformants were selected based on hygromycin resistance and the stable lines were obtained in 3^rd^ or 4^th^ generation. Two independent overexpression lines driven by the *35S* promoter, namely *35S:BBX13#1* and *35S:BBX13#2*, were used for the experiments. The *co 35S:BBX13#1, ft-10 35S:BBX13#1,* and *35S:CO-YFP 35S:BBX13#1* lines were generated by crossing *35S:BBX13#1* with the respective single mutant lines.

### GUS Histochemical assay

The full-length promoter of *BBX13* (∼1.5kb) was amplified from the Col-0 genomic DNA using Gateway-compatible primers (**Table S1**) and was cloned into *pDONR207* using BP cloning method. The *proBBX13:GUS* construct was generated by performing an LR reaction between the *pGWB3* vector and *pDONR207-proBBX13* construct. The *proBBX13:GUS* construct was then used for transforming Col-0 plants, and independent stable transformants were selected using hygromycin resistance. Histochemical GUS staining was performed as previously described by Weigel and Glazebrook, (2002).

### Flowering time and rosette leaf quantification

After three days of stratification, the plates were grown under long-day or short-day conditions in a Percival growth chamber. The 10-day-old seedlings were transferred to the soil and maintained in a growth room under similar conditions with a light fluence of ∼100 μmol m^−2^ s^−1^. The flowering time and number of rosette leaves were recorded when the plants started bolting and the bolt reaches up to ∼1cm long.

### Yeast two-hybrid (Y2H) assay

Yeast two-hybrid assays were performed according to the Clontech manual as described previously (Ravindran et al., 2021). Full-length CDS of *CONSTANS, BBX13, BBX13-N-terminal, BBX13-Middle domain, BBX13-C-terminal*, *BBX28* and *FT* were cloned into the *pDONR207* vector and then subcloned into the *pDEST-GADT7* or *pDEST-GBKT7* vectors to generate the bait and prey constructs. The plasmids were then transformed into the AH109 yeast strains by PEG transfection. Positive colonies (yeast cells containing both bait and prey) were selected on yeast (Synthetic Drop-out) SD-Leu-Trp plates and further dropped onto SD-Leu-Trp, SD-Leu-Trp-His, SD-Leu-Trp-His-Ade, and SD-Leu-Trp-His + 3-AT (3-Amino-1,2,4-triazole). The 3-AT concentrations are mentioned wherever used.

### Bimolecular fluorescence complementation assay

The BiFC protocol was followed as suggested by Gampala et al., (2007). The full-length CDS of *BBX13* and *CO* was cloned into the *pCL112* and *pCL113* vectors, respectively, by LR gateway cloning from the *pDONR* constructs. The full-length CDS of the positive and negative interaction controls *BBX28* and *FT* were cloned into the *pCL112* vector, by LR gateway cloning from the *pDONR* constructs. The *pCL112* vector has an N-terminal region of YFP, and the *pCL113* vector has a C-terminal region of YFP. The plasmids were then infiltrated into the tobacco leaves using Agrobacterium-mediated infiltration. After two days of keeping the plants in the dark, the leaves were cut and imaged under a confocal microscope for fluorescence.

### RNA isolation, cDNA synthesis, and Quantitative real-time PCR

All RNA isolation experiments were performed using the Takara RNAiso Plus, following the manufacturer’s instructions. cDNA was synthesized using 1µg of total RNA using the Takara PrimeScript 1st strand cDNA Synthesis Kit, following the manufacturer’s protocol. Real-time PCR was performed in a LightCycler®96 (Roche, www.roche.com) machine using TB Green® Premix EX Taq™ II SYBR Green dye (TaKaRa, www.takarabio.com) following the protocol using 1μl of cDNA per reaction as a template. *UBIQUITIN10* and *GAPC2* genes were used as internal controls.

### Electrophoretic mobility shift assay (EMSA)

The *TF-HIS-BBX13* (BBX13 fused to the Trigger Factor chaperone and a HIS tag) construct was generated by cloning the *BBX13* CDS into the *pCOLD-TF* vector using KpnI and HindIII as restriction sites. The construct was transformed into BL21 cells, and the transformed cells were induced with IPTG (0.5mM) at 28°C. TF-HIS (HIS tag along with a Trigger Factor chaperone) and TF-HIS-BBX13 were purified using the Ni-NTA beads. The *CORE2* motif of the *FT* promoter was synthesized as the DNA fragment. DNA-protein complex was detected with the Chemiluminescent Nucleic Acid Detection Module (Thermo Fisher Scientific), using the manufacturer’s instructions. The EMSA protocol was performed as previously described (Job et al., 2018).

### Chromatin Immunoprecipitation (ChIP) qPCR

The ChIP assay was performed as described previously (Saleh et al., 2008; Lau and Bergmann, 2015; Komar et al., 2016). First, 10-day-old LDs grown seedlings were harvested, and the tissue was crosslinked by fixing in 1% formaldehyde. Chromatin was then isolated using a series of extraction buffers as described by Saleh et al., (2008), followed by two cycles of sonication with 30s on/30s off at 4°C. The sonicated chromatin-protein complex was allowed to bind to the bead-bound antibody. Anti-GFP (GF28R, Invitrogen) and mouse IgG control (10400C, Invitrogen) antibodies were used. The bead-bound chromatin was then processed for reverse crosslinking, and the beads were removed from the solution using a magnetic rack, followed by DNA purification. 10% of the total sonicated chromatin was used as the input DNA control. Enrichment was checked by qPCR using primers specific to the promoter/protein-binding region. Percentage input methods were used to analyze the qPCR data (Haring et al., 2007; Yadukrishnan et al., 2020).

### Protoplast isolation and Dual-Luciferase assay

Protoplasts were isolated from Col-0 leaves following the protocol described by Yoo et al., (2007). We used *BBX13* and *CO* cloned in the modified *pCAMBIA1300* vector under the *35S* promoter as effectors and a ∼2kb promoter fragment of *FT* cloned into the *pGreenII-0800-LUC* vector as the reporter construct. We examined how BBX13 affects the CO-mediated activation of *FT* transcription. The vectors were then transfected into protoplasts using PEG transfection. Luciferase activity was analyzed using the Promega Kit (E1910) as per protocol after overnight incubation of transfected protoplast. Renilla luciferase activity was used as an internal control (Ravindran et al., 2021).

### Statistical analysis

GraphPad Prism V8.0 was used to prepare graphs and perform statistical analyses. All the analyses are described in the respective figure legends.

### Accession numbers

*BBX13* (AT1G28050), *CO* (AT5G15840), *FT* (AT1G65480), *SOC1* (AT2G45660), *TSF* (AT4G20370), *AP1* (AT1G69120), *LFY* (AT5G61850), *FLC* (AT5G10140), and *BBX28* (AT4G27310).

## Results

### *BBX13/COL15* expression is spatially and temporally regulated

BBX13/COL15 is one of the few B-box family members in Arabidopsis that remain elusive for their physiological function. To understand the expression pattern of *BBX13*, we generated *proBBX13:GUS* reporter lines, and GUS histochemical staining was performed to visualize the activity of the *BBX13* promoter in Arabidopsis tissues. We grew the seedlings in Murashige and Skoog (MS) plates for 10 days and then transferred them to the soil under long days (LDs - 16h light/ 8h dark). The expression of *BBX13* showed a gradual increase in seedlings from 3-10 days after stratification (DAS), primarily in the vasculature, including the veins of the cotyledons and the true leaves (**Figure 1A**). GUS staining was also detected in the shoot apex region of seedlings and flowers of the adult plants (**Figure 1A**). Next, we quantified the relative transcript levels of *BBX13* in the various tissues of Col-0 at different growth stages. The transcript levels of *BBX13* were high in mature rosette and cauline leaves, buds, and flowers (**Figure 1B**). Further, we quantified the expression of *BBX13* at different times of the day following the Zeitgeber time (ZT) and found that *BBX13* follows a distinct diel expression pattern (**Figure 1C**). The transcript levels peak 8-12 hours after dawn, after which the levels decrease (**Figure 1C**). Taken together, our data indicate that *BBX13* expression is spatially and temporally regulated.

**Figure 1.**
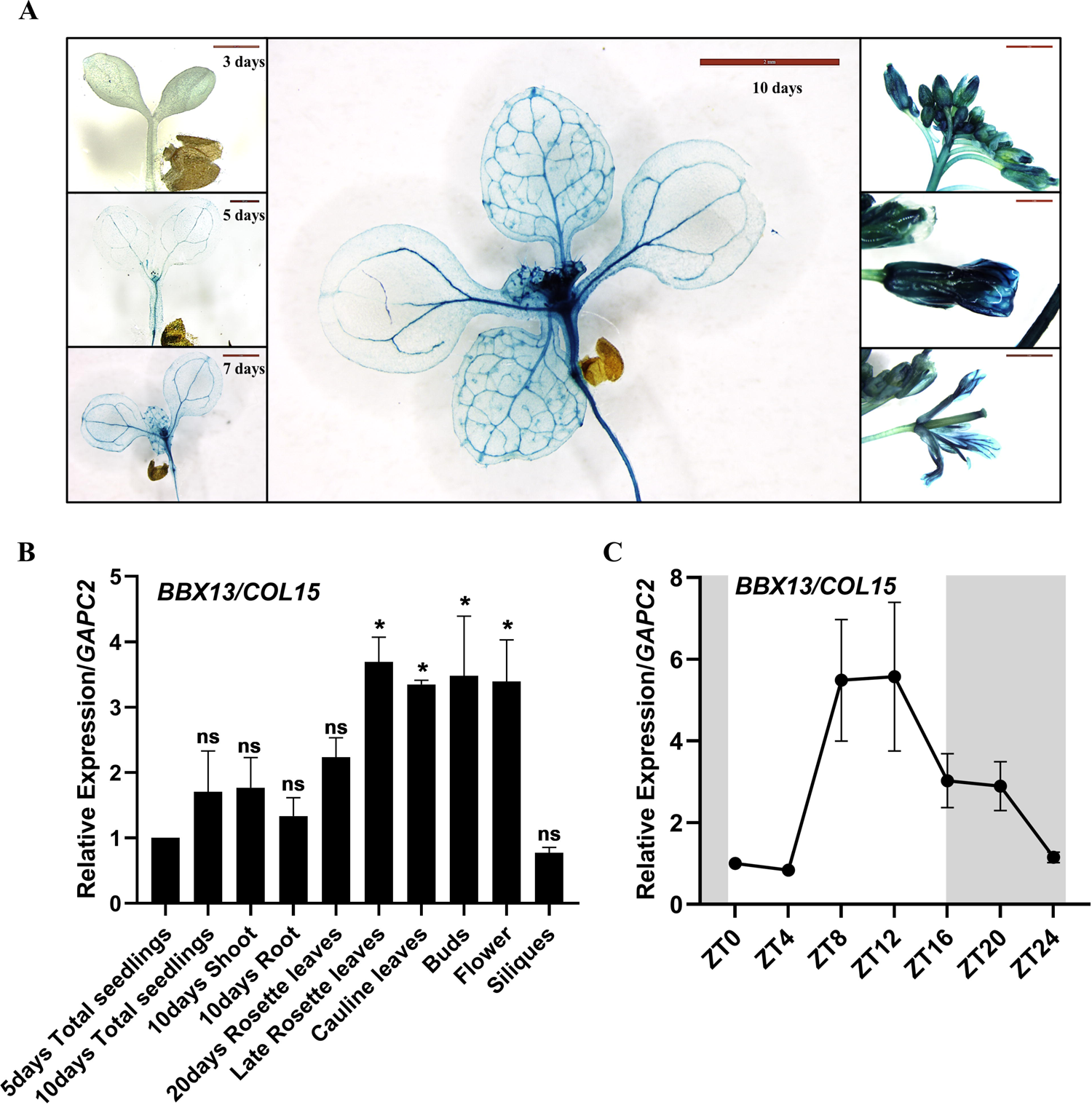
Spatio-temporal and developmental regulation of *BBX13/COL15* expression (A) Histochemical staining of *proBBX13:GUS* lines in the 3 days, 5 days, 7 days, and 10 days-old seedlings showing the activity mainly in the veins, leaves, and shoot apex region. Seedlings were grown for 10 days in MS media and then transferred to the soil under long-day conditions (LDs - 16h/8h). The right side panels show GUS staining, indicating the expression of *BBX13* in the buds and flowers in adult *proBBX13:GUS* plants. (B) RT-qPCR results show the relative transcript levels of *BBX13* at ZT12 in the indicated tissue samples across developmental stages from wild-type (Col-0) plants grown under LDs. *GAPC2* was used as the internal control. Data are mean ± SEM, n=2. Asterisks represent statistically significant differences as determined by one-way ANOVA followed by Dunnett’s multiple comparison test (*, *P* < 0.05; ns, not significant). (C) Temporal expression pattern of *BBX13* in the wild-type seedlings. RNA was isolated from 10-day-old seedlings grown in the LDs. Samples were collected at a 4-hour interval following the Zeitgeber time (ZT). The grey part indicates the 8h night period while the 16h white part is daytime. *GAPC2* was used as the internal control. Data are mean ± SEM, n=3.

### BBX13/COL15 negatively regulates flowering time in Arabidopsis under inductive photoperiods

The expression pattern of *BBX13* in the veins of leaves and cotyledons is remarkably similar to the expression pattern of *CO* and *FT*, which are key regulators of flowering time (An et al., 2004; Kumar et al., 2012). To understand if BBX13/COL15 plays a role in the regulation of flowering, we quantified the flowering time in the mutants and overexpressors of *BBX13*. We obtained two T-DNA-insertion mutants, *bbx13-1* and *bbx13-2,* from the stock center and also generated two independent lines that constitutively overexpress *BBX13/COL15*, namely *35S:BBX13#1* and *35S:BBX13#2.* Our qRT-PCR analysis indicated 80-fold and 90-fold upregulation in *BBX13* expression in *35S:BBX13#1* and *35S:BBX13#2,* while some residual expression was detected in the mutants (**Figure S1A,B).** *bbx13-1* and *bbx13-2* fail to produce full-length transcripts of *BBX13* (**Figure S1A,C).** To further verify if the qPCR expression values obtained for the *bbx13* mutants are background values, we performed a semi-qPCR using two different primer pairs that span the region amplified by the qPCR primers. The forward primers (CDS_F and M_F) bind to different regions of exon1, while the reverse primer (M_R) binds to exon3, which can amplify any transcripts arising from this region (**Figure S1A**). We detected the bands (BBX13-Δ1 and BBX13-Δ2) in the Col-0 sample (Figure S1D,E) using these primers. We detected a faint band for *BBX13-Δ2* in *bbx13-1,* however the absence of a distinct band in the *bbx13* mutants probably suggests the absence of any robustly expressed truncated transcripts in the T-DNA lines (**Figure S1E**). Under inductive long-day conditions (LDs - 16h light/8h dark), *bbx13* mutants flowered ∼2-3 days earlier than Col-0, whereas the *BBX13*-overexpressors exhibited a severe delay of more than 15 days in flowering time. We also quantified the number of rosette leaves at bolting. Under long days, the mutants exhibited significantly lower number of leaves, whereas the lines overexpressing BBX13 produced ∼14 more rosette leaves than Col-0 (**Figure 2A-C**). Expression levels of *BBX13* and the extent of delay in flowering time seemed to have a positive correlation as *35S:BBX13#2,* which possessed higher *BBX13* levels, flowered 3 days later with 1-2 extra rosette leaves compared to *35S:BBX13#1* (**Figure 2A-C, Figure S1B**). To examine whether the role of BBX13 in the regulation of flowering time is photoperiod-dependent, we quantified the flowering time in the above genotypes under short-day conditions (SDs - 8h light/16h dark).

**Figure 2.**
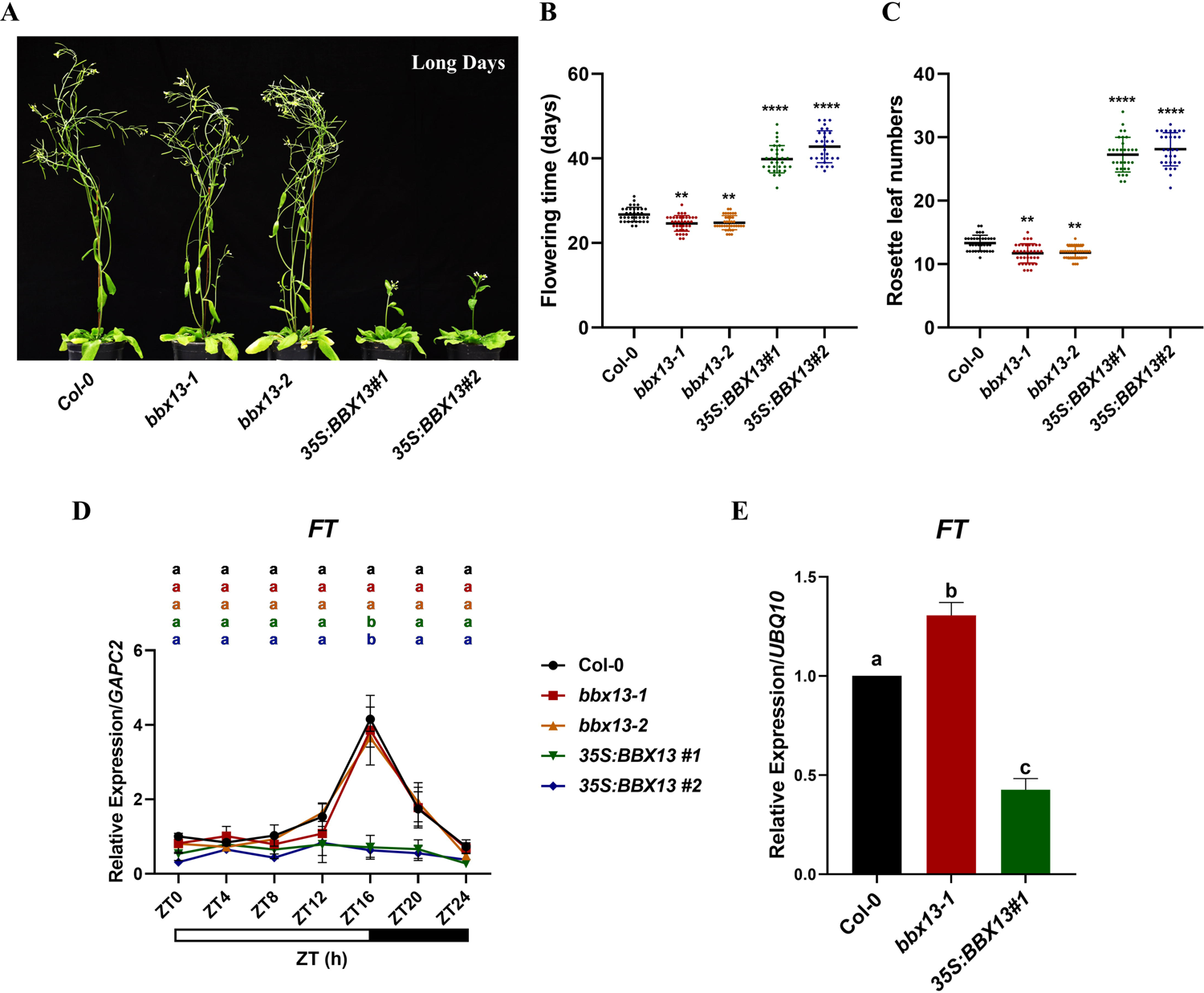
BBX13 negatively regulates flowering time under long-day conditions by suppressing *FT* transcript levels (A) Representative images of 45 days-old plants of Col-0*, bbx13-1, bbx13-2, 35S:BBX13#1* and *#2* grown under long-day conditions (LDs - 16h/8h). (B) Flowering time quantified under LDs. The days to flower were calculated as the number of days taken from seed germination to bolting. The scatter dot plot represents the mean ± sd with the total number of plants ≥30. (C) Rosette leaf numbers quantified under LDs. The number of rosette leaves was counted once the plants started bolting. The scatter dot plot represents the mean ± sd with the total number of plants ≥ 30. Asterisks represent statistically significant differences (* *P* < 0.05, ** *P* < 0.01, *** *P* < 0.001, **** *P* < 0.0001, ns-not significant) as determined by one-way ANOVA followed by Dunnett’s multiple comparisons test. (D) Relative expression of *FT* in 10-day-old seedlings grown under long days. RNA was isolated from seedlings collected at every 4h Zeitgeber (ZT). *GAPC2* was used as the internal control. Data are mean ± SEM, n=2. The white box from ZT0-ZT16 below the graphs indicates the 16h daytime while the black box from ZT16-ZT24 indicates the 8h night time. (E) Relative expression of *FT* in the shoot region of 20-day-old seedlings grown under long days. Seedlings for RNA were collected at ZT16. *UBQ10* was used as the internal control. Data are mean ± SEM, n=2. Letters denote the statistical groups obtained using one-way ANOVA, followed by Tukey’s multiple comparisons test (P < 0.05).

Unlike long-day conditions, all genotypes flowered at similar time under short-day conditions (**Figure S2A-B**). We have also quantified the rosette leaf numbers under short days, while the mutants bolted with a similar number of leaves compared to Col-0, the overexpressors bolted with almost 10% less leaves (**Figure S2C)**. Since we could notice some developmental aberrations in the leaf size and shape in the overexpression lines, possibly contributing to the differences in the leaf numbers, we decided to use only “days to flowering” as the quantifying parameter for our subsequent experiments. Together, our results indicate that BBX13/COL15 negatively regulates flowering time in a photoperiod-dependent manner.

### BBX13/COL15 negatively regulates flowering through transcriptional repression of *FT* by interacting with CO

Next, we investigated the effect of loss-of-function and overexpression of *BBX13/COL15* on the expression of the major photoperiodic flowering time regulator genes. We examined the expression pattern of *CO, FT, SOC1, TSF (TWIN SISTER OF FT), AP1, LFY,* and *FLC (FLOWERING LOCUS C)* in 10-day-old seedlings grown under long days. The *FT* transcript levels in 10-day-old seedlings were drastically decreased in the overexpression lines compared to the wild-type and the mutants, specifically at ZT16 (**Figure 2D)**. In shoots of 20-day-old seedlings, *FT* transcript levels were significantly upregulated in the *bbx13-1* compared to the wild-type, which might account for the early flowering phenotype of *bbx13* mutants (**Figure 2E**). The *CO* expression in the mutants did not show a significant difference compared to wild-type (**Figure S3A)**. We could detect a significant downregulation in *CO* expression in *35S:BBX13#1* at ZT20 (**Figure S3A)**. We did not observe any significant changes in the transcript levels of *FLC,* which is known to regulate flowering through the autonomous and vernalization pathways (**Figure S3B**). We found that the expression levels of *TSF, AP1* and *LFY,* were significantly decreased in the overexpression lines during ZT16 to ZT24, while no significant change in the *SOC1* expression levels was observed among the genotypes (**Figure S3C-F)**. These results suggest that BBX13 negatively regulates the flowering time by repressing the transcript levels of major positive regulators of the photoperiodic pathway. CO promotes flowering by transcriptionally activating *FT* (Samach et al., 2000). Since *BBX13* and *CO* share a similar spatial expression pattern in the veins and leaves, we asked whether BBX13 modulates the CO-mediated activation of *FT.* Previous reports suggest that a number of proteins modulate the function of CO through protein-protein interaction (Song et al., 2020; Takagi et al., 2023). Moreover, BBX proteins are known to physically interact with other members in the family to regulate each other’s function (Yadav et al., 2020a; Shin et al., 2023). Therefore, we examined whether BBX13 interacts with CO. To check the interaction in-planta, we performed a bimolecular fluorescence complementation (BiFC) assay by co-transforming different combinations of split YFP constructs into *Nicotiana benthamiana* leaves. YFP fluorescent signals were observed only when nYFP-BBX13 and cYFP-CO were co-transformed, validating the interaction between the two proteins (**Figure 3A**). To validate this interaction, we have used BBX28 as a positive control (Liu et al., 2020) and FT as a negative control for the interaction with CO (**Figure 3A**). We further performed a yeast two-hybrid assay with BBX13 and CO fused to activation and binding domains, respectively and confirmed that they physically interact (**Figure 3B**). As mentioned earlier, this interaction was also validated by using BBX28 and FT as positive and negative controls (**Figure 3B**). To map the CO-interacting domain of BBX13, we divided full-length BBX13 into three fragments (N, M, and C) containing the B-box domains, the middle domain, and the CCT domain, respectively (**Figure 3C)**. We cloned them separately into the binding domain (BD) containing vector and investigated their interaction with activation domain (AD) tagged CO in yeast (**Figure 3D**). Among the three parts, only the N-terminal fragment of BBX13 containing the B-box domain interacted with full-length CO (**Figure 3C-D**). The middle fragment was not used in the assay as it showed autoactivation. Together, these results show that BBX13 physically interacts with CO through its B-box domain. Although the expression pattern of *BBX13* in the veins of the leaves is similar to that of *FT*, we could not detect physical interaction between BBX13 and FT in our yeast two-hybrid experiments (**Figure S4**).

**Figure 3.**
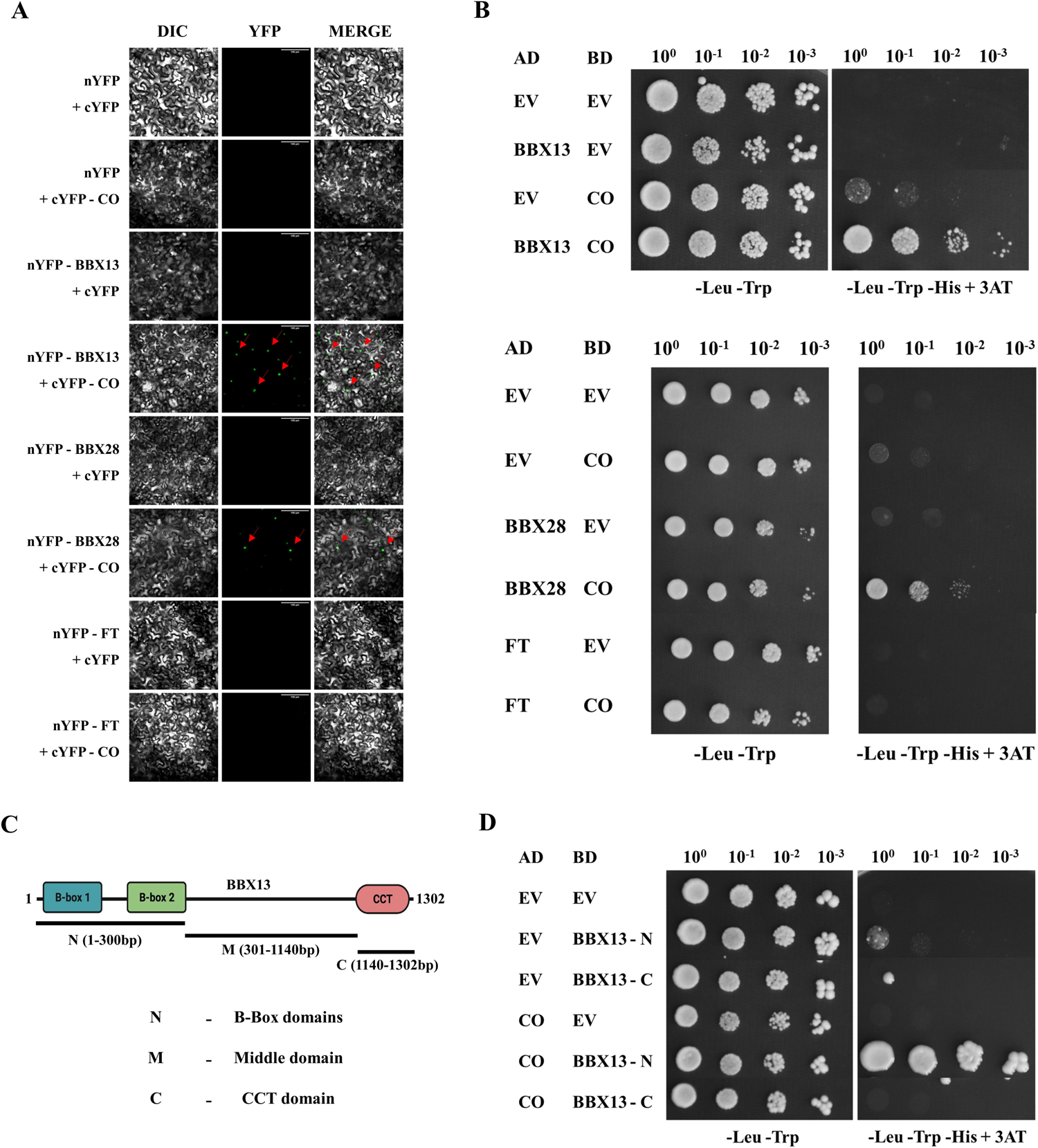
BBX13 physically interacts with CO. (A) Bimolecular fluorescence complementation (BiFC) assay showing in-planta interaction of BBX13 with CO. nYFP and cYFP refer to the N-terminal and C-terminal of Yellow Fluorescent Protein, respectively. The labels on the left side of the panels indicate the combinations with which the *Nicotiana benthamiana* leaves were infiltrated. The labels above the panels indicate the channels that the images were captured. The red arrows indicate some of the signals captured from the respective channels. BBX28-CO and FT-CO combinations were used as positive and negative controls, respectively. The scale bar indicates 100µm. (B) Yeast two-hybrid assay showing the interaction of BBX13 with CO. AD and BD refers to the activation domain and binding domain of GAL4 in *pDEST-GADT7* and *pDEST-GBKT7* vectors, respectively. EV indicates an empty vector. -Leu-Trp indicates the double drop-out media without leucine and tryptophan. -Leu-Trp-His + 3AT indicates the triple drop-out media without leucine, tryptophan, and histidine, supplemented with 10 mM 3AT (3-Amino-1,2,4-triazole). The numbers mentioned above the panels are serial dilutions for drop-test. BBX28-CO and FT-CO combinations were used as positive and negative controls for the interaction, respectively. (C) The schematic shows how BBX13 has been divided to make N, M, and C domains for the Y2H assay. The exact base position where the truncations were made is given in parentheses. (D) Yeast two-hybrid assay showing the interaction of BBX13-N terminal region with CO. CO is in the activation domain (AD), while BBX13-N and BBX13-C are in the binding domain (BD). The BBX13-M domain shows autoactivation and is not represented.

### *BBX13* genetically interacts with *CO* in the photoperiod pathway

To investigate the genetic interaction between *BBX13* and *CO*, we generated the double mutant *co bbx13-1* and quantified the flowering time under long-day conditions. The *co bbx13-1* plants exhibited a substantial delay in flowering time similar to *co*, suggesting that *co* is epistatic to *bbx13-1* (**Figure 4A-B)**. In addition, we crossed *35S:BBX13#1* with *35S:CO-YFP* to generate a double-overexpression line and compared its flowering time with the parental lines and the wild type under long days. We checked the transcript levels of *CO* and *BBX13* in the double overexpressor line using RT-qPCR to verify whether silencing of any of the genes had occurred. Both the genes show similar levels of expression in the double OE lines compared to the single overexpressor parental lines, suggesting that they are not silenced in the double OE line (**Figure S6A-B**). The *35S:CO-YFP 35S:BBX13#1* double overexpressor line flowered earlier than the Col-0, almost similar but slightly delayed compared to *35S:CO-YFP* (**Figure 4C-D)**.

**Figure 4.**
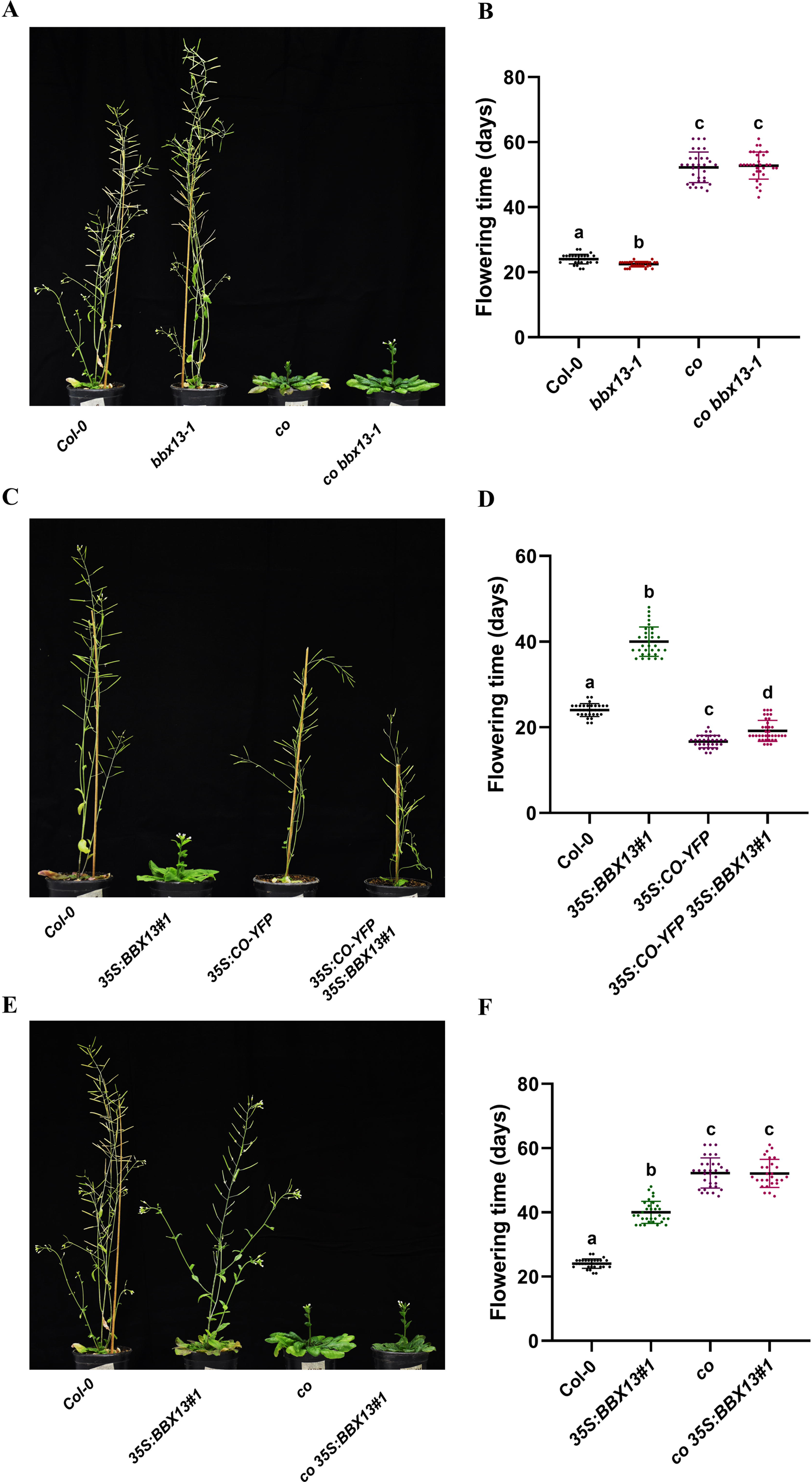
*BBX13* and *CO* genetically interact to regulate flowering Representative images of (A, E) 53 days-old plants and (C) 41 days-old plants grown under long-day conditions (LDs - 16h/8h). (B, D, F) Flowering time quantified under LDs. The days to flower were calculated as the number of days taken from seed germination to bolting. Scatter dot plots represent the mean ± sd with the total number of plants ≥30. Letters denote the statistical groups obtained using Brown-Forsythe and Welch ANOVA tests, followed by Games-Howell’s multiple comparisons test (P < 0.05).

Furthermore, we crossed *co* with *35S:BBX13#1* to generate *co 35S:BBX13#1* line which flowered at a similar time to *co* (**Figure 4E-F)**. These results suggest that *BBX13* regulates photoperiodic flowering in a *CO*-dependent manner. We also generated the double mutants *ft-10 bbx13-1* and *ft-10 35S:BBX13#1*, and both lines showed a delay in flowering similar to that of *ft-10* (**Figure S5A-D**). Altogether, our results confirm that BBX13 physically and genetically interacts with CO and acts through the *CO-FT* module to delay flowering in Arabidopsis.

### BBX13 disrupts CO-mediated activation of *FT*

BBX proteins are known to regulate the transcription of their target genes by directly binding to the upstream regulatory regions (Xu et al., 2016; Song et al., 2020; Yadav et al., 2020a; Cao et al., 2023; Singh and Datta, 2023). Since *FT* expression is altered in *BBX13*-overexpressors, we performed an electrophoretic mobility shift assay (EMSA) to investigate if BBX13 binds to the *FT* promoter. Previous reports have shown that the strongest activation of *FT* by CO is known to be via its binding on the *CORE2* motif, while *CORE1* binding exhibits lesser activity (Tiwari et al., 2010). In order to examine if BBX13 can directly bind to the *CORE2* motif, we performed an EMSA. Our results indicate that BBX13 directly binds to the *CORE2* motif present in the promoter of *FT* (**Figure 5A-B**). Since CO is known to bind to the same site, BBX13 may compete with CO to occupy this region on the *FT* promoter (Kumimoto et al., 2008; Tiwari et al., 2010). As BBX13 and CO interact physically, we asked if BBX13 modulates the binding of CO to the *FT* promoter. To test this, we compared the enrichment of CO on the *FT* promoter in *35S:CO-YFP* and *35S:CO-YFP 35S:BBX13#1* seedlings by performing a chromatin immunoprecipitation (ChIP)-qPCR. We amplified the P1 fragment on the *FT* promoter, containing the *CORE1* and *CORE2* motifs which act as potential binding sites for CO (**Figure 5C**). The enrichment of CO on the promoter of *FT* was decreased in *35S:CO-YFP 35S:BBX13#1* as compared to *35S:CO-YFP,* suggesting that BBX13 negatively regulates the binding of CO on the *FT* promoter (**Figure 5C-D)**. This reduced binding possibly results in the decreased transcript levels of *FT* in the *35S:CO-YFP 35S:BBX13#1* compared to *35S:CO-YFP* (**Figure S6C**). To further examine the ability of BBX13 to inhibit the CO-mediated transcriptional activation of *FT*, we performed a dual-luciferase assay in wild-type Arabidopsis protoplasts. We co-transfected *proFT:LUC* reporter with *35S:CO* and *35S:BBX13* effectors in different combinations (**Figure 5E-F**). The presence of *35S:CO* caused strong induction of *proFT:LUC*, whereas only *35S:BBX13* had no significant effect. When both the effectors *35S:CO* and *35S:BBX13* were present together at equimolar concentrations, the luciferase signals were decreased compared to that by *35S:CO*. When the relative concentration of *35S:BBX13* was increased further, we could observe a strong decrease in the luciferase signals induced by *35S:CO*. These results indicate that increasing concentrations of BBX13 quantitatively inhibit the CO-mediated activation of *FT*.

**Figure 5.**
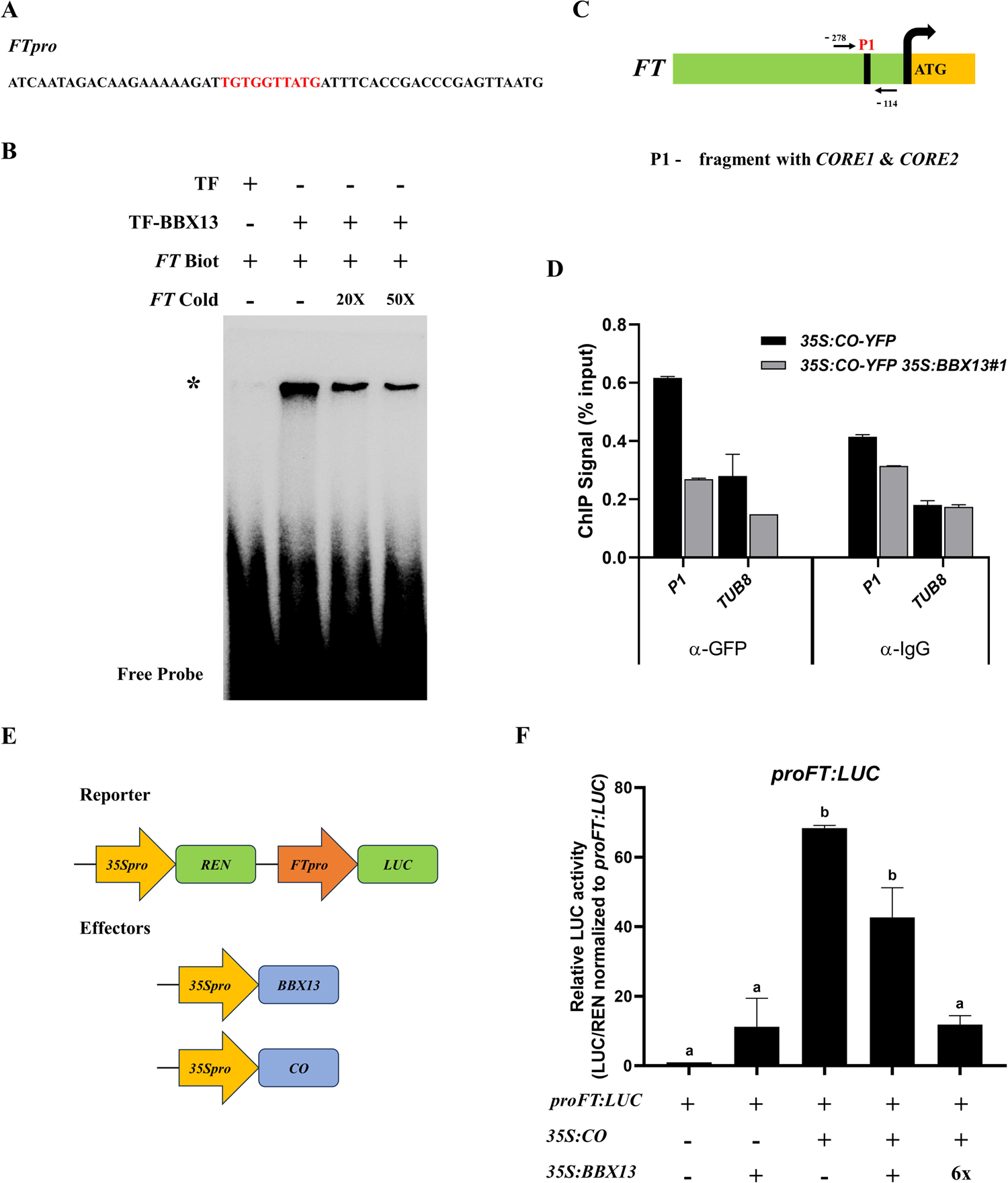
BBX13 binds to *FTpro* and inhibits CO-mediated transcriptional activation of *FT* (A) The sequence of the promoter region of *FT* used as a probe for the electrophoretic mobility shift assay (EMSA). The red highlighted region is the *CORE2* motif ‘TGTG(N2-3)ATG’. (B) EMSA shows the binding of TF-BBX13 (BBX13 fused to the ‘Trigger Factor’ chaperone and HIS tag) to the *CORE2* motif in the promoter of *FT*. The biotinylated probe with *CORE2* motif is indicated as *FT* Biot. *FT* Cold indicates the non-biotinylated probe. + and – represent presence and absence. Competition for the labelled probe was performed by adding 20x and 50x unlabelled cold probe. The asterisk indicates the shifted band of BBX13 bound to the *FT* promoter. (C) Schematic of the *FT* promoter. The green box indicates the putative promoter region, and the yellow box represents the genomic region. The black bent arrow represents the transcription start site. The black rectangle inside the green box represents the P1 fragment on the *FT* promoter, containing the *CORE1* and *CORE2* motifs, which act as potential binding sites for CO. Regular black arrows below or above the promoter mark the primer binding sites and the number below them represents the nucleotide position of the primers used for ChIP-qPCR. (D) ChIP-qPCR analysis determining CO binding onto the *FT* promoter in seedlings of *35S:CO-YFP* and *35S:CO-YFP 35S:BBX13#1* grown under long-day conditions for 10 days. Seedlings were harvested at ZT14. DNA-protein complexes were immunoprecipitated using anti-GFP. Immunoprecipitants using anti-IgG antibody were used for negative control. *TUB8* was used as a non-target negative control in the qPCR. (E) Schematic of the constructs used in the transient assay in *Arabidopsis* protoplast. The reporter construct used was generated on the *pGreenII-0800 Luc* vector expressing *35S promoter*-driven Renilla Luciferase (REN) and *FTpro*-driven Luciferase (LUC). *BBX13* and *CO* cloned under the constitutive *35S* promoter were used as effectors. (F) Relative luciferase assay showing the effect of BBX13 on CO-mediated activation of the *FT* promoter. The effectors and *proFT:LUC* were co-transfected into the Arabidopsis protoplasts. The relative luciferase activity was measured after overnight incubation. Data are mean ± SEM with n=2. Letters denote the statistical groups obtained using one-way ANOVA, followed by Tukey’s multiple comparisons test (*P* < 0.05).

## Discussion

Flowering is a major developmental transition in the life-cycle of a plant, that leads to reproductive development, mating, and seed production. To ensure that flowering happens under environmental conditions that support further reproductive development, plants have evolved a complex network of regulators coordinating the external and internal cues (Koornneef et al., 1998; Perrella et al., 2020). Decades of research led to the identification of the CO-FT module as the central regulatory hub in the photoperiodic flowering pathway (Srikanth and Schmid, 2011; Takagi et al., 2023). Importantly, the homologs of key regulators of flowering time that were initially identified in Arabidopsis were found to be functionally conserved in other crop plants, while some other genes have acquired diverse functions (Blümel et al., 2015). Allelic variations in flowering time genes, especially those in the photoperiod and vernalization pathways, have facilitated the adaptation of crop plants to different geographical regions (Lin et al., 2021). Early flowering is a useful trait in regions where various environmental factors constrain the durations of growing seasons, whereas delayed flowering is beneficial in the case of bioenergy and fodder crops as it enhances the vegetative biomass (Nuñez et al., 2017).

BBX proteins are known to play roles in a wide range of processes such as germination, seedling establishment, cotyledon opening and greening, biotic and abiotic stress, hormone-signaling, photomorphogenesis and thermo-morphogenesis, UV signaling, shade avoidance, flowering time regulation, etc. (Vaishak et al., 2019; Song et al., 2020; Yadav et al., 2020a; Yadav et al., 2020b; Cao et al., 2023; Singh et al., 2023). In this study, we identified that a previously uncharacterized group-II BBX protein, BBX13/COL15 negatively regulates flowering time in Arabidopsis under long-day conditions. *BBX13* exhibits spatial expression pattern similar to *CO* and *FT* (**Figure 1**) (An et al., 2004; Kumar et al., 2012). CO is known to bind to the *FT* promoter to positively regulate *FT* transcription (Wenkel et al., 2006; Tiwari et al., 2010). BBX13/COL15, when overexpressed, causes significant reduction in *FT* expression leading to a severe late flowering phenotype in long days (**Figure 2; Figure S1,2,3**). We found that BBX13 physically interacts with CO (**Figure 3; Figure S4**). Genetic analyses showed that *BBX13* act upstream to *CO* and *FT* in the flowering pathway (**Figure 4; Figure S5**). We also show the direct binding of BBX13 to the *FT* promoter by EMSA. Our ChIP data indicates that BBX13 reduces the in vivo binding of CO on the *FT* promoter (**Figure 5A-D**). Through luciferase assay, we found that BBX13 inhibits the CO-mediated transcriptional activation of *FT* (**Figure 5E-F)**

Multiple BBX proteins have been found to impart both positive and negative effects on flowering time in Arabidopsis and other crops (Crocco and Botto, 2013; Song et al., 2020; Talar and Kiełbowicz-Matuk, 2021; Bandara et al., 2022; Cao et al., 2023). BBX10, BBX17, BBX19, BBX28, and BBX30/31 suppress flowering by physically interacting with CO, leading to the inhibition of its activity (Wang et al., 2014; Graeff et al., 2016; Liu et al., 2020; Song et al., 2020; Xu et al., 2022). In most of these cases, B-box domain, particularly the B-box motif 1, was found to be indispensable for their physical interaction with CO. In agreement, we observed that BBX13/COL15 interacts with CO via its N-terminal region that harbors the B-box domain. Together, these studies corroborate that the inhibition of CO function through physical interaction is perhaps the most commonly adopted mechanism by the BBX proteins to suppress flowering in Arabidopsis. However, it is important to note that the evidence for such mechanism in other plant species remains obscure in spite of the identification of several BBX proteins as suppressors of flowering, especially in Rice and Chrysanthemum (Talar and Kiełbowicz-Matuk, 2021; Cao et al., 2023).

Since BBX proteins are known to form heterodimers within the family, it is tempting to speculate that BBX13/COL15 and the other BBX proteins may heterodimerize to interact with CO or may act as parts of a larger BBX multiprotein complex that regulate flowering time by modulating the function of CO. For example, BBX4 has shown a direct binding to the proximal region of *FT* promoter through the interaction with BBX32, thus regulating flowering time (Tripathi et al., 2016). Concordantly, a recent study employed yeast two-hybrid screening to show extensive heterodimerization between BBX proteins, which includes mutual interactions between BBX13, BBX19, BBX28, and BBX31 (Shin et al., 2023).

It is possible that different BBX proteins feed diverse external and internal information to CO-FT module to optimize the functioning of the reproductive switch. For example, *BBX28* and its close homolog *BBX29* are induced by low ambient temperatures and play a role in determining flowering time under these conditions (Wang et al., 2021). *BBX31* is known to be regulated by external signals like UV-B and hormonal signals like ABA (Yadav et al., 2019; Singh and Datta, 2023). It has been shown recently that chloroplast-derived retrograde signals induce the expression of *BBX14/15/16* via GOLDEN2-LIKE1/2 (GLK1/2) and the physical interaction of BBX14/15/16 with CO inhibits the ability of CO to activate *FT* and thus to promote flowering (Susila et al., 2023). In addition, the expression of multiple BBX genes, including those regulating the flowering time, was proposed to be under the control of the circadian clock (Ledger et al., 2001; Kumagai et al., 2008; Tripathi et al., 2016). BBX19, a repressor of flowering, interacts with PSEUDO RESPONSE REGULATOR proteins (PRRs) and facilitates the suppression of morning-phased clock genes, including the central clock-associated transcription factor CIRCADIAN CLOCK ASSOCIATED 1 (CCA1) (Yuan et al., 2021). Interestingly, a combinatorial approach using ChIP-seq and genome-wide expression profiling has identified *BBX13* as one of the potential direct targets of CCA1, indicating that *BBX13* could possibly be a clock-regulated gene (Kamioka et al., 2016). This is further supported by our observation that *BBX13* follows a diel expression pattern that peaks towards the middle of the day and comes down in the evening (**Figure 1C**). It can be speculated that a direct repression by CCA1 in the morning could be responsible for the evening-phased expression pattern of *BBX13*. The rhythmicity of *BBX13* may vary with species since *CmBBX13*, a likely ortholog of *AtBBX13* in Chrysanthemum, showed a completely arhythmic expression irrespective of the photoperiod (Ping et al., 2019). Our data also suggest that BBX13 has a more prominent effect on the expression of *FT* at adult vegetative stages compared to the juvenile stage, indicating a possible modulation of the function of BBX13 by the developmental age of the plant (**Figure 2D-E**). In the future, it will be interesting to explore the regulation of *BBX13* by various environmental and developmental signals and how they influence its function in flowering time regulation.

In summary, our study unveils the role of a previously uncharacterized B-box protein BBX13/COL15 in controlling the flowering time in Arabidopsis (**Figure 6**). We show that BBX13/COL15 modulates the photoperiodic flowering pathway by physically interacting with CO and attenuating the CO-mediated transcriptional activation of key flowering-inducing genes like *FT*. Exploring the possibility of modulating BBX13/COL15 and associated regulatory modules to tailor flowering time in crop plants can be a promising direction for future research.

**Figure 6.**
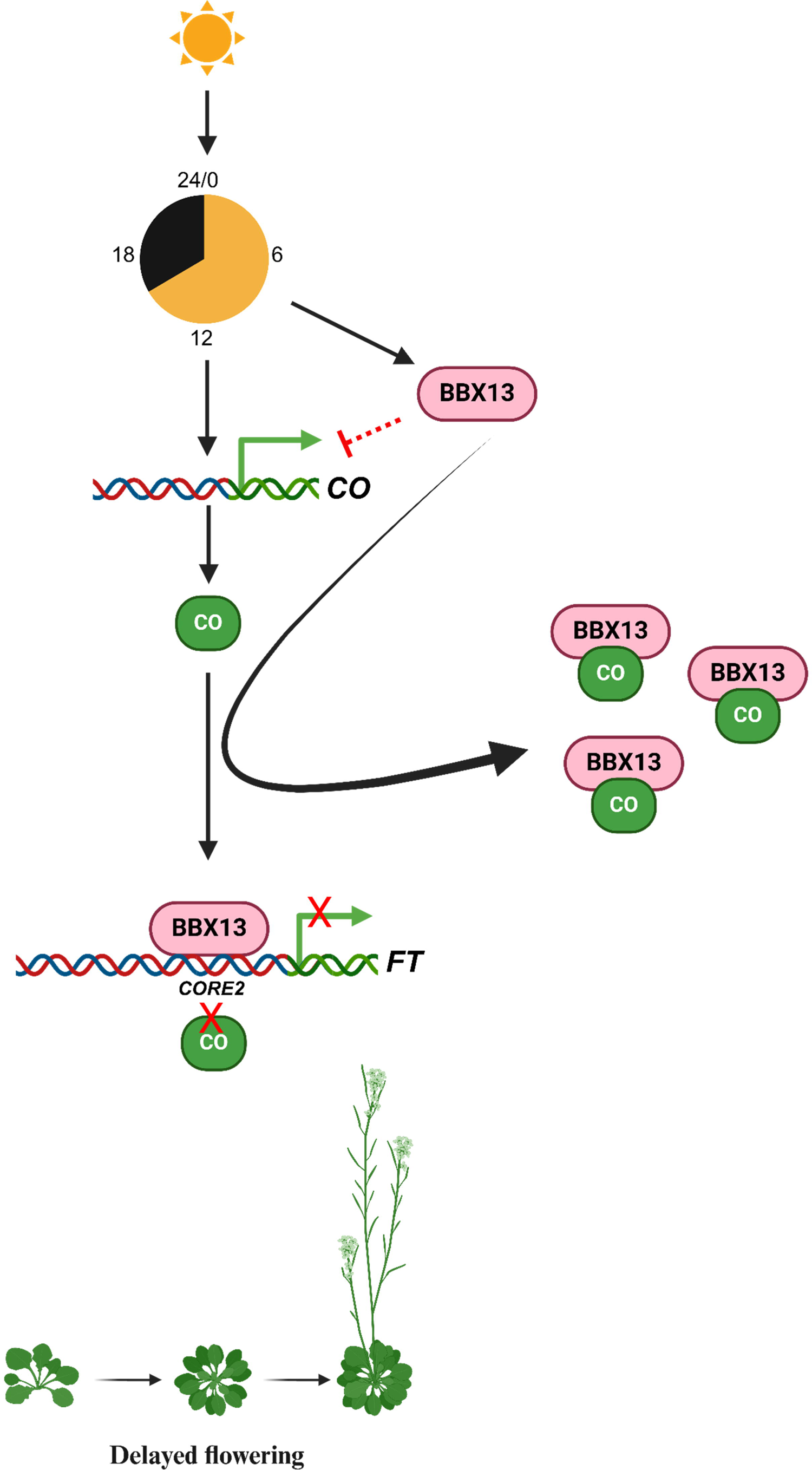
Proposed model showing the role of BBX13 in regulating photoperiodic flowering in Arabidopsis. Under long-day conditions, BBX13 negatively regulates flowering by modulating the CO-FT module in Arabidopsis. BBX13 downregulates the transcription of *CO*. BBX13 physically interacts with CO and sequesters it to inhibit the CO-mediated transcriptional activation of *FT*. In addition, BBX13 directly binds to the *CORE2* motif on the *FT* promoter, a site where CO can also bind, to inhibit *FT* activation and delay flowering.

## ACKNOWLEDGEMENTS

The authors thank Prithiv Raj for his initial help in genotyping double mutants. They thank Prof. Utpal Nath of IISc Bengaluru for kindly providing the *35S:CO-YFP* seeds. They thank Shubhi Dwivedi for the BBX28 constructs and Arpan Mukherjee, Debojyoti Kar, Venkateswara Rao and other PCDB lab members for valuable comments.

## AUTHOR CONTRIBUTIONS

SD, PVR and PY conceived the study and designed the experiments. PVR performed all the experiments; PY helped in generating the overexpressor lines; AS helped in generating double mutants and performing Y2H. PVR and SD analyzed the data. PVR, PY and SD wrote the manuscript. SD supervised the overall study. All authors revised the final manuscript.

## Supplemental Data

**Figure S1.** Sites of T-DNA insertions in the *bbx13* mutants and the expression levels of *BBX13* in mutants and overexpressors

**Figure S2.** BBX13 does not alter flowering time under short-day conditions

**Figure S3.** BBX13 negatively regulates transcript levels of CO and other flowering regulator genes but exerts no significant effects on FLC expression

**Figure S4.** BBX13 does not show physical interaction with FT in yeast

**Figure S5.** FT acts epistatic to BBX13

**Figure S6.** Transcript levels of BBX13, CO, and FT in lines overexpressing BBX13, CO, and both

**Table S1:** List of primers used in this study

