## Supplemental File for "The B-box protein BBX13/COL15 suppresses photoperiodic flowering by attenuating the action of CONSTANS in Arabidopsis"

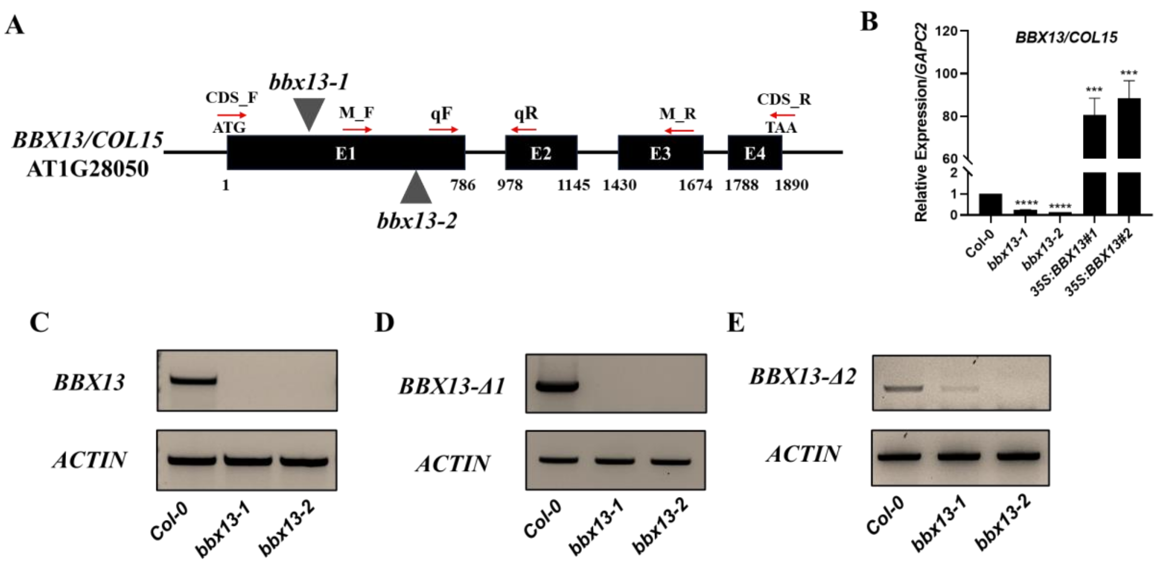


**Figure S1. Sites of T-DNA insertions in the *bbx13* mutants and the expression levels of *BBX13* in mutants and overexpressors**

(A) Gene model of *BBX13/COL15* with the site of T-DNA insertion in *bbx13-1* and *bbx13-2* indicated by grey arrowheads. ATG represents the start codon, and TAA represents the stop codon. E1-E4 black boxes indicate the four exons in the *BBX13,* and the lines between the black boxes indicate introns. (B) RT-qPCR shows the transcript levels of *BBX13* in the mutants and overexpressors compared to Col-0. The *BBX13* qPCR primers are labeled qF and qR with red horizontal arrows over the gene model. Numerals after # indicate the independent lines of overexpressors. RNA was isolated from 10-day-old seedlings grown under long-day conditions (LDs - 16h/8h). *GAPC2* was used as the internal control. Data are mean ± SEM, n=3. Asterisks represent statistically significant differences (*** $P < 0.001$, **** $P < 0.0001$, ns-not significant) as determined by one-way ANOVA followed by Dunnett's multiple comparisons test. (C) Semiquantitative RT-PCR of *BBX13* CDS shows that the *bbx13-1* and *bbx13-2* mutants fail to produce full-length transcripts of *BBX13*. *BBX13* CDS forward (CDS_F) and reverse (CDS_R) primers were used to amplify the full-length transcript of 1302 base pairs. (D) *BBX13-Δ1* represents semiquantitative RT-PCR of truncated *BBX13* amplified using *BBX13* CDS forward (CDS_F) and M-domain reverse primers (M_R). The expected size of *BBX13-Δ1* is 1140 base pairs. (E) *BBX13-Δ2* represents semiquantitative RT-PCR of truncated *BBX13* amplified using M-domain forward (M_F) and reverse primers (M_R). The expected size of *BBX13-Δ2* is 840 base pairs. The primers used are labeled with red horizontal arrows over the gene model. *ACTIN* was used as the loading control. RNA was isolated from 20 days-old seedlings grown in LDs.

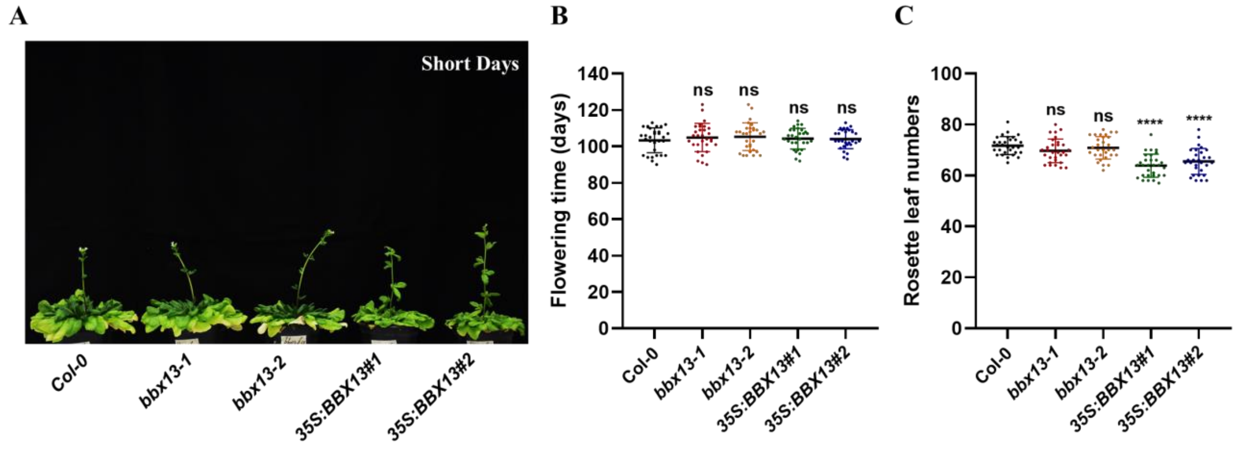


**Figure S2. BBX13 does not alter flowering time under short-day conditions**

(A) Representative images of 110 days-old plants of Col-0*, bbx13-1, bbx13-2, 35S:BBX13#1* and *#2* grown under short-day conditions (SDs - 8h/16h). (B) Flowering time quantified under SDs. The days to flower were calculated as the number of days taken from seed germination to bolting. The scatter dot plot represents the mean ± sd with the total number of plants ≥30. (C) Rosette leaf numbers quantified under SDs. The number of rosette leaves was counted once the plants started bolting. The scatter dot plot represents the mean ± sd with the total number of plants ≥ 30. Asterisks represent statistically significant differences (* $P < 0.05$, ** $P < 0.01$, *** $P < 0.001$, **** $P < 0.0001$, ns-not significant) as determined by one-way ANOVA followed by Dunnett's multiple comparisons test.

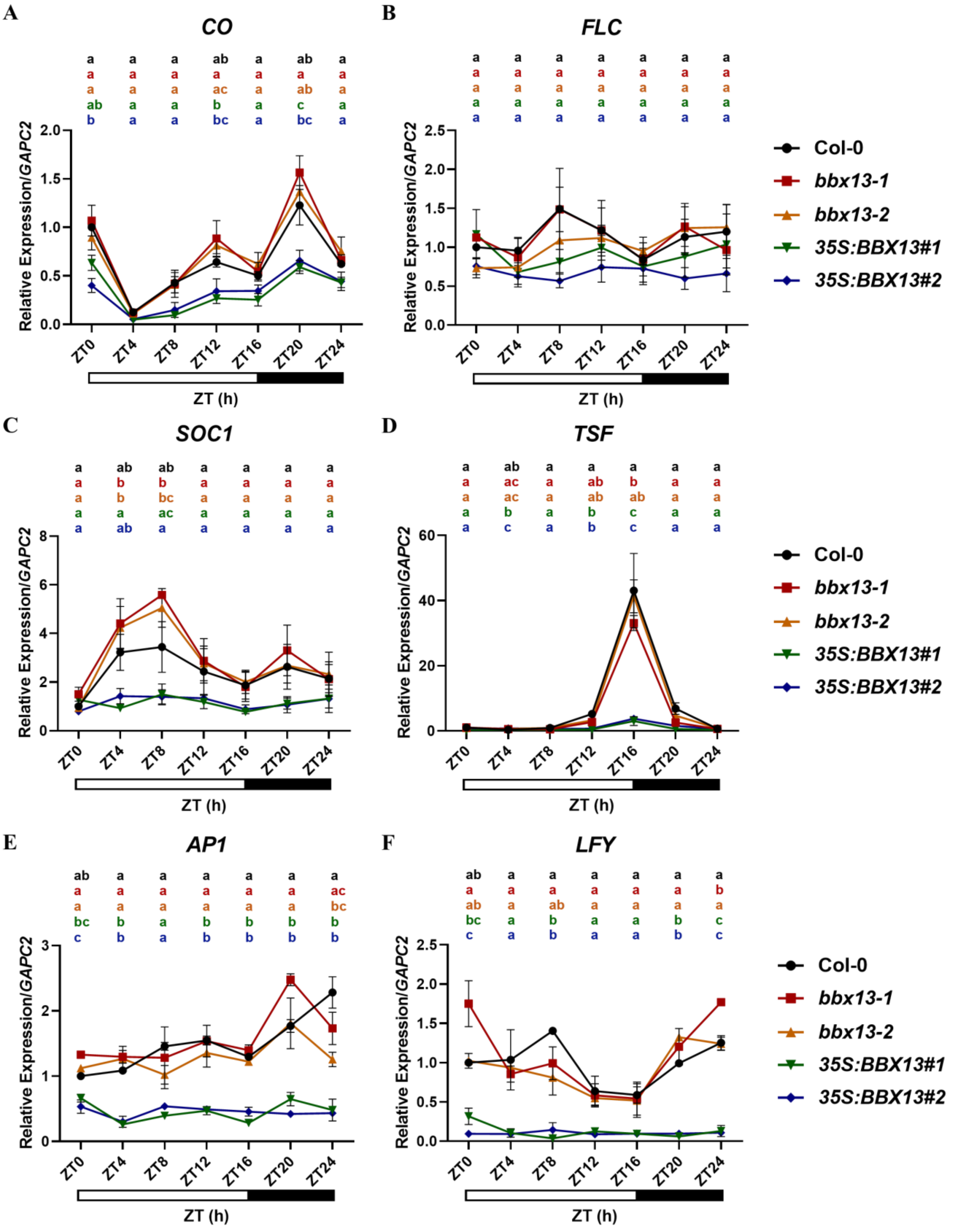

A
CO
B
FLC
C
SOC1
D
TSF
E
AP1
F
LFY
Relative Expression/GAPC2
ZT0
ZT4
ZT8
ZT12
ZT16
ZT20
ZT24
ZT (h)
Col-0
bbx13-1
bbx13-2
35S:BBX13#1
35S:BBX13#2

**Figure S3. BBX13 negatively regulates transcript levels of *CO* and other flowering regulator genes but exerts no significant effects on *FLC* expression**

(A-F) Relative expression of the major flowering-pathway genes *CO, FLC, SOC1, TSF, AP1,* and *LFY* in 10-day-old seedlings grown under long days. RNA was isolated from seedlings collected at every 4h Zeitgeber (ZT). *GAPC2* was used as the internal control. Data are mean ± SEM, n=2. The white box from ZT0-ZT16 below the graphs indicates the 16h daytime while the black box from ZT16-ZT24 indicates the 8h night time. Letters denote the statistical groups obtained using one-way ANOVA, followed by Tukey's multiple comparisons test ($P < 0.05$).

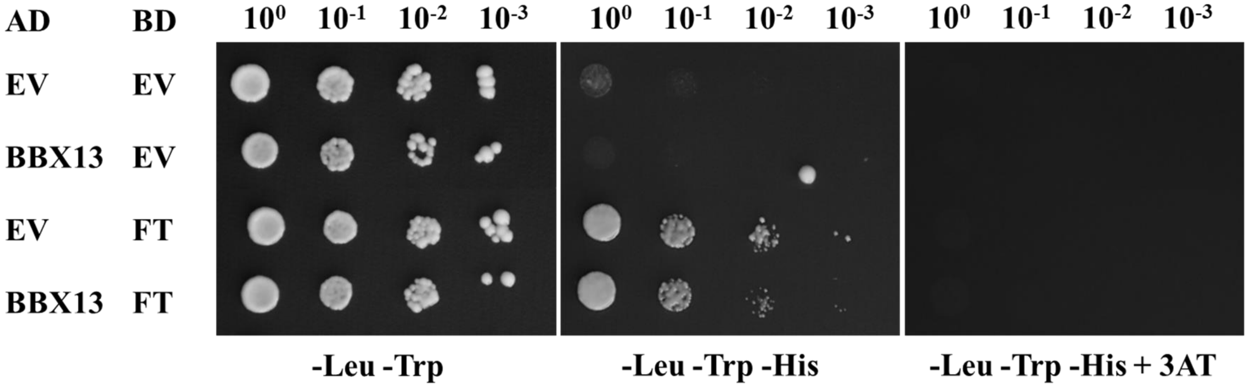


**Figure S4. BBX13 does not show physical interaction with FT in yeast**

The yeast two-hybrid assay shows that BBX13 does not interact with FT. AD and BD refer to the activation and binding domains of GAL4 in *pDEST-GADT7* and *pDEST-GBKT7* vectors, respectively. EV indicates an empty vector. -Leu-Trp indicates the double dropout media without leucine and tryptophan. -Leu-Trp-His indicates the triple dropout media without leucine, tryptophan, and histidine. -Leu-Trp-His + 3AT indicates the triple dropout media without leucine, tryptophan, and histidine, supplemented with 10 mM 3AT (3-Amino-1,2,4-triazole). The numbers mentioned above the panels are serial dilutions for drop-test.

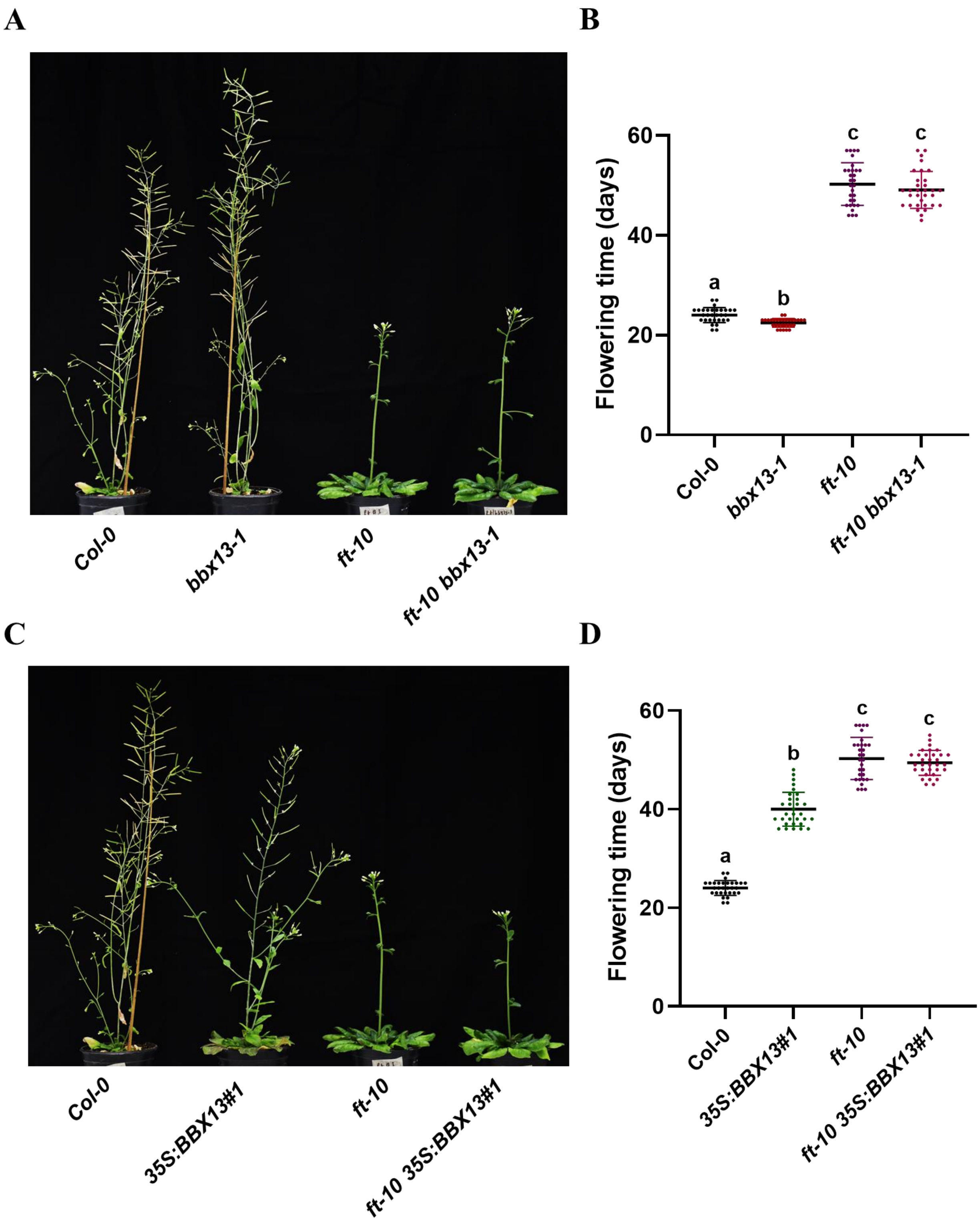
A
Col-0
bbx13-1
ft-10
ft-10 bbx13-1
B
Flowering time (days)
60
40
20
0
a
b
c
c
Col-0
bbx13-1
ft-10
ft-10 bbx13-1
C
Col-0
35S:BBX13#1
ft-10
ft-10 35S:BBX13#1
D
Flowering time (days)
60
40
20
0
a
b
c
c
Col-0
35S:BBX13#1
ft-10
ft-10 35S:BBX13#1

**Figure S5. *FT* acts epistatic to *BBX13***

Representative images of (A, C) 53 days-old plants grown under long-day conditions (LDs - 16h/8h). (B, D) Flowering time quantified under LDs. The days to flower was calculated once the plants started bolting from the day of germination. Scatter dot plots represent the mean ± sd with the total number of plants ≥30. Letters denote the statistical groups obtained using Brown-Forsythe and Welch ANOVA tests, followed by Games-Howell's multiple comparisons test (P < 0.05).

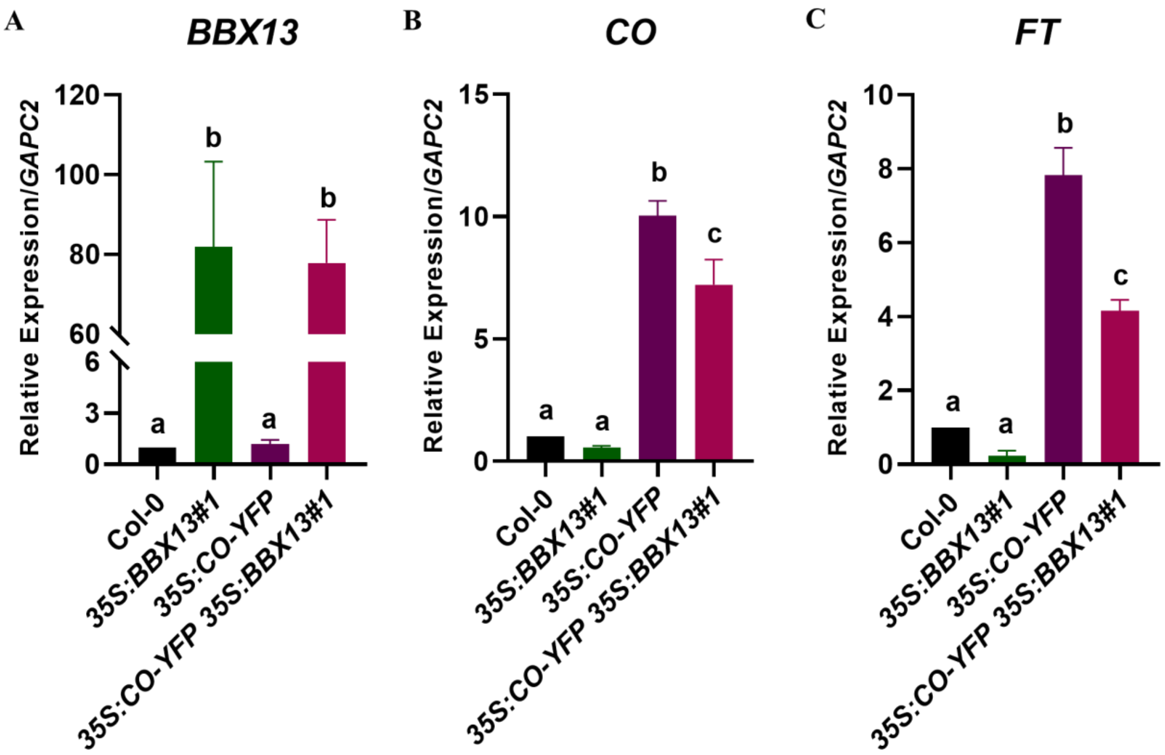


**Figure S6. Transcript levels of *BBX13, CO,* and *FT* in lines overexpressing *BBX13*, *CO,* and both**

(A-C) Relative expression of *BBX13, CO,* and *FT* in 10-day-old seedlings grown in long days. Seedlings for RNA were collected at ZT12. *GAPC2* was used as the internal control. Data are mean ± SEM, n=3. Letters denote the statistical groups obtained using one-way ANOVA, followed by Tukey's multiple comparisons test (*P* < 0.05).

**Table S1: List of primers used in this study**

| Primer Name | Primer Sequence (5' - 3') |
|---|---|
| *bbx13-1 LP* | AAACCTGAGAACTCTGGCCAC |
| *bbx13-1 RP* | GAACAATGGCGACTCTCTCAG |
| *bbx13-2 LP* | ACGGGAGATGACAACGTATTG |
| *bbx13-2 RP* | TACGGCGGTTTTGTTTTGTAG |
| *LBb1.3* | ATTTTGCCGATTTCGGAAC |
| *co-sail LP* | AAGCTGTTGTGACACATGCTG |
| *co-sail RP* | CCCCTTCTTTCAGATACCAGC |
| *LB2* | GCTTCCTATTATATCTTCCCAAATTACCAATACA |
| *ft-GK LP* | GGTGGAGAAGACCTCAGGAAC |
| *ft-GK RP* | TTTTGGGAGACAAATTGATGC |
| *o8409* | ATATTGACCATCATACTCATTGC |
| *GUS Internal F* | CTTACGCTGAAGAGATGCTC |
| *GUS Internal R* | CGCGATCAAAGACGCG |
| *35Spro END F* | CGCAAGACCCTTCCTCTAT |
| *ACT2 F* | ACCCAAAGGCCAACAGAGAG |
| *ACT2 R* | TGAACGATTCCTGGACCTGC |
| *proBBX13_attB1* | GGGGACAAGTTTGTACAAAAAAGCAGGCTGGAATTTGCTTCT GATTCGCCG |
| *proBBX13_attB2* | GGGGACCACTTTGTACAAGAAAGCTGGGTGAACTGACTCTC TCTCAAAATCC |
| *BBX13_attB1 (CDS_F)* | GGGGACAAGTTTGTACAAAAAAGCAGGCTGGATGAGTAGTT CGGAGAGAG |
| *BBX13_attB2 (CDS_R)* | GGGGACCACTTTGTACAAGAAAGCTGGGTGTTAAGGGTAAG GAGCTTCAC |
| *CO_attB1* | GGGGACAAGTTTGTACAAAAAAGCAGGCTGGATGTTGAAAC AAGAGAGTAACGAC |

| | |
|---|---|
| *CO_attB2* | GGGGACCACTTTGTACAAGAAAGCTGGGTGTCAGAATGAAGGAACAATCC |
| *BBX13_N_attB1* | GGGGACAAGTTTGTACAAAAAAGCAGGCTGGATGAGTAGTTCGGAGAGAG |
| *BBX13_N_attB2* | GGGGACCACTTTGTACAAGAAAGCTGGGTGACAACCGGAAAAACCTTC |
| *BBX13_M_attB1 (M_F)* | GGGGACAAGTTTGTACAAAAAAGCAGGCTGGCCATCGGCGTTGGAGCT |
| *BBX13_M_attB2 (M_R)* | GGGGACCACTTTGTACAAGAAAGCTGGGTGCCGCTCCAGATCAGCCTTAG |
| *BBX13_C_attB1* | GGGGACAAGTTTGTACAAAAAAGCAGGCTGGCTGGCTCAGAACAGAGG |
| *BBX13_C_attB2* | GGGGACCACTTTGTACAAGAAAGCTGGGTGTTAAGGGTAAGGAGCTTCA |
| *BBX28_attB1* | GGGGACAAGTTTGTACAAAAAAGCAGGCTTAATGGGGAAGAAGTGTGATTTATG |
| *BBX28_attB2* | GGGGACCACTTTGTACAAGAAAGCTGGGTCTTAAACAACAACCGTTGATTTAAACG |
| *FT_attB1* | GGGGACAAGTTTGTACAAAAAAGCAGGCTGGATGTCTATAAATATAAGAGACCCTCTTATAGTAAGC |
| *FT_attB2* | GGGGACCACTTTGTACAAGAAAGCTGGGTGCTAAAGTCTTCTTCCTCCGCA |
| *proFT F Xho1* | CCGCTCGAGAGTGGCAGATACGTTAAATTTTATAA |
| *proFT R Hind111* | CCCAAGCTTCTTTGATCTTGAACAAACAGGT |
| *BBX13 qF* | TGAGGAGATCAATGGTGGCG |
| *BBX13 qR* | ACTCGTATCCTCAGGTCCCC |
| *CO qF* | CAACAGCTTCACACCCAAGAACG |
| *CO qR* | TTGCAGGGTCAGGTTGTTGCTC |
| *FT qF* | GCTACAACTGGAACAACCTTTGGC |
| *FT qR* | TGAATTCCTGCAGTGGGACTTGG |

| | |
|---|---|
| *SOC1 qF* | TTCGCCAGCTCCAATATGCAAG |
| *SOC1 qR* | TGCTGACTCGATCCTTAGTATGCC |
| *TSF qF* | GTGCCGAGTCCAAGCAAC |
| *TSF qR* | CGATGAATTCCCGAGGGGG |
| *AP1 qF* | AATATGCCTCCCCCTCTGC |
| *AP1 qR* | CGGGTTCAAGAGTCAGTTCGA |
| *LFY qF* | TGATGCTCTCTCCCAAGAAGGG |
| *LFY qR* | TCAGTCTGGTCTTGTTGCTGCAC |
| *FLC qF* | TGTTCAACTGGAGGAACACCTTG |
| *FLC qR* | AGCTTCAACATGAGTTCGGTCTTC |
| *UBQ10 qF* | GGCCTTGTATAATCCCTGATGAATAAG |
| *UBQ10 qR* | AAAGAGATAACAGGAACGGAAACATAGT |
| *GAPC2 qF* | TTGGTGACAACAGGTCAAGCA |
| *GAPC2 qR* | AAACTTGTCGCTCAATGCAATC |
| *BBX13 KpnI F* | CGGGGTACCATGAGTAGTTCGGAGAGAGTACC |
| *BBX13 HindIII R* | CCCAAGCTTAGGGTAAGGAGCTTCAC |
| *FT-CORE2 EMSA F* | ATCAATAGACAAGAAAAAGATTGTGGTTATGATTTCACCGACCCGAGTTAATG |
| *FT-CORE2 EMSA R* | CATTAACTCGGGTCGGTGAAATCATAACCACAATCTTTTTCTTGTCTATTGAT |
| *FT P1 ChIP qF* | GGTTTGGAATACCACAAACAGAAA |
| *FT P1 ChIP qR* | ACTGTTTCGGATTTGCATTAACTCG |
| *TUB8 ChIP qF* | CCGTTTCAAATTCTCTCTCTC |
| *TUB8 ChIP qR* | CAAACTACTTCCCAGAACTTAGC |
